# DISTRIBUTION OF THE ENDANGERED GIANT KANGAROO RAT, *DIPODOMYS INGENS*, ON THE NAVAL PETROLEUM RESERVES, CALIFORNIA

**DOI:** 10.1101/062240

**Authors:** Thomas P. O’Farrell, Nancy E. Mathews, Patrick M. McCue, Mary S. Kelly

## Abstract

Burrows of the endangered giant kangaroo rat, *Dipodomys ingens*, were found on the U.S. Department of Energy’s Naval Petroleum Reserves in Kern County, California. The majority of burrows (63%) were found in areas of low relief (4.3 ± 0.5°, n=32) on valley floors but 28% were unexpectedly found on low relief areas (6.4 ± 0.8°, n=39) in the uplands. Burrows were not distributed in proportion to the arealextend of the soil series, but were concentrated in deep alluvial sandy loams. Well pads were the most frequently observed (73% of 211) human disturbance in the vicinity of burrows, but they were the most distant (150± 61m, n=150). Dirt roads were observed closer to burrows (50 ± 10 m, n=28) but less frequently (4%). Only 8 burrows were found in the vicinity of proposed petroleum developments but no projects had to me modified to avoid negatively affecting the species.

RESUMEN.—Madrigueras de las ratas canguro, *Dipodomys ingens,* a riesgo de extinsioń, fueron localizadas en el Departmento de Energía, Reservas Navales de Petroleo en el Condado de Kern, California. La mayoria de las madrigueras (63%) se encuentran en arias de bajo relieve (4.3 ± 0.5°, n=32) en valles, pero el 28% fueron inesperadamente localizadas en arias de bajo relieve (6.4 ± 0.8°, n=39) en terrenos elevados. Las madriguera no se distribuyen en proporcion de acuerdo a la variedad de terrenos, sino concentradas en las lomas aluviales arenosas. Los disturbios humanos observados con más frecuencia (73% of 211) en la vecindad de las madrigueras son los almohadillas de pozos petroleros pero se encontran amayor distancia (150 ± 61m, n=150). Caminos de tierra fueron encontrados cerca de las madrigueras (50 ± 10m, n=28) pero con menos frecuenciz (4%). Unicamente ocho madrigueras fueron localizadas en la vecindad de los propuestos desarroyos petroleros pero ningun proyecto se debe modificar para evitar efectos negativos que pueden afectar a la especie.

---

The giant kangaroo rat, *Dipodomys ingens,* historically inhabited a narrow strip of arid steppe habitat that included seven California counties along the western side of the San Joaquin Valley from about Los Banos, south to Taft, and on the Carrizo Plain, Elkhorn Plain, and in the Cuyama Valley. Within this range populations were widely scattered and only locally abundant (Grinnell 1932). After concluding his field work Grinnell (1932) became aware of a soils map of his study area (Nelson et al. 1921) and observed that the areas mapped as Panoche fine sandy loam, “…include most of those in which *D. ingens* has been found.” and that, “…future field work is urgently suggested in the tracing and verification or disproving of this seemingly strong correlation.” When much of the historical distribution was resurveyed the species occupied habitats in a wider range of soil types (Williams 1992, Williams et al. 1995).

Because much of the original habitat of the giant kangaroo rat was lost to agricultural and energy developments, and urbanization, the United States Fish and Wildlife Service (FWS) listed it as endangered under the Endangered Species Act (ESA) on 5 January 1987 (United States Department of the Interior 1987).

Our objective was to determine whether the species occurred on the U. S. Department of Energy’s (DOE) Naval Petroleum Reserves (NPR-1, NPR-2) in the south-central part of the San Joaquin Valley. If it did, we wanted to obtain information that would help DOE develop and implement a conservation plan for the species in compliance with ESA. An ancillary goal was to test Grinnell’s (1932) hypothesis that the distribution of giant kangaroo rats was correlated with the distribution of specific silt loams. If it were confirmed, the information could be used to define critical habitat and focus conservation efforts there.

## Methods

### Study Areas

NPR-1 and NPR-2 are located 42 km southwest of Bakersfield, Kern County, California (Fig. 1). NPR-1 consists of 191 km^2^ and is located on the northwestern border of NPR-2 which includes 122 km^2^. Both were established in 1912 to provide a source of petroleum for the United States Navy. They were under the jurisdiction of the Navy until 1977 when they were transferred to the United States Department of Energy. NPR-1 was sold to Occidental Petroleum Corporation in 1998, and NPR-2 was transferred to the United States Department of the Interior in August, 2005.

**Fig. 1.**
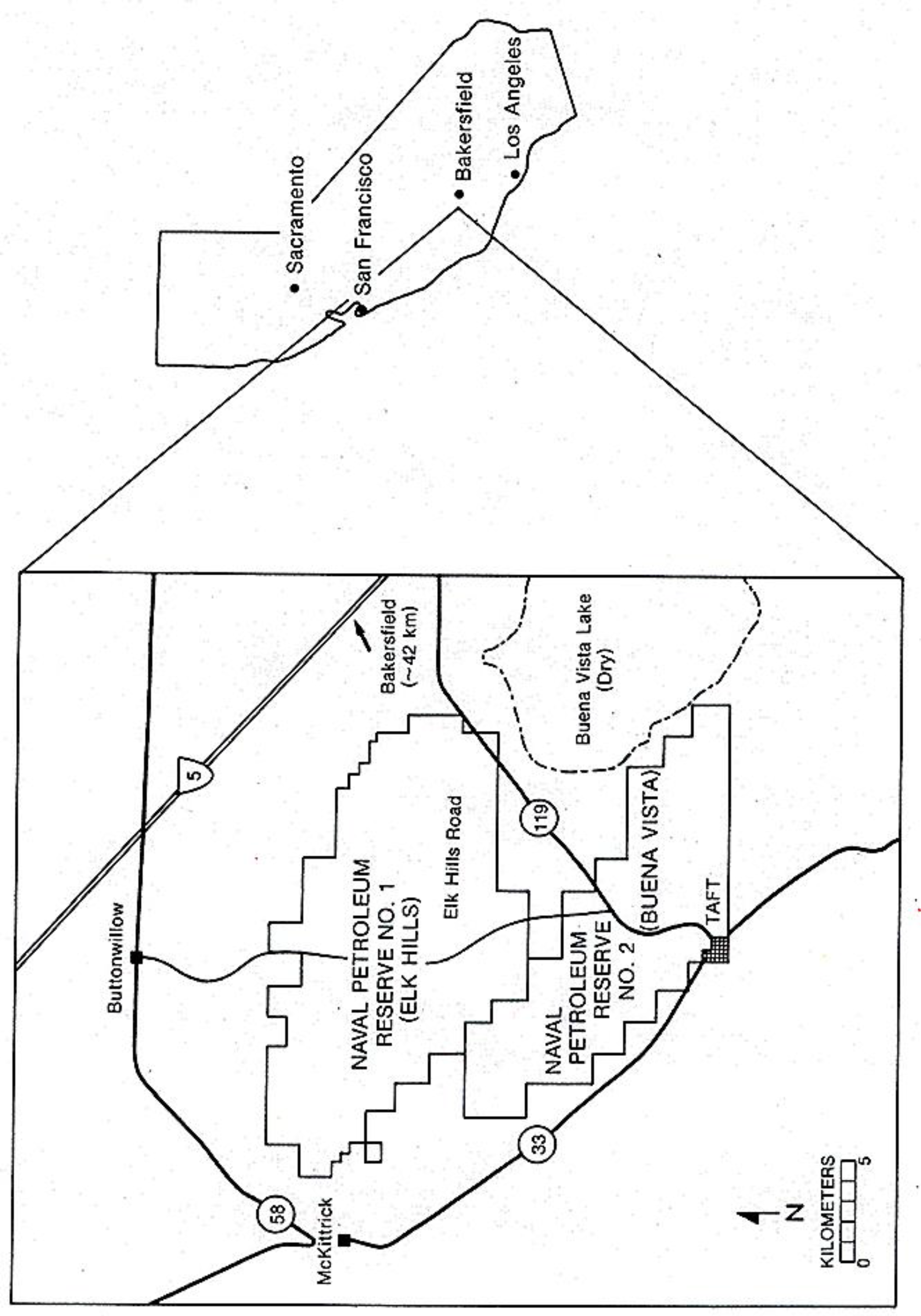
Location of the Naval Petroleum Reserves, Kern County, California.

NPR-1 is bisected by the Elk Hills which are low foothills of the Temblor Range that extend southeastward into the San Joaquin Valley. The Buena Vista Hills form a northwest-southeast crest along the length of NPR-2 and are roughly parallel with the Elk Hills to the north. Together these two sets of hills form a region of dissected terrain rising from the floor of the south-central edge of the San Joaquin Valley just east of, but disjunct from, the steep Temblor Range. Elevations range between 88-473 m above sea level on NPR-1 and between 94-393 m on NPR-2.

The Mediterranean climate consists of hot, dry summers and cool, damp winters. The mean maximum temperature in summer is 35° C: the mean minimum temperature is 18° C. The mean maximum temperature in winter is 17° C: the mean minimum temperature is 5° C. Nearly all precipitation, averaging 114-152 mm annually, occurs during the winter months of November-March.

Soils are residual or alluvial, and derived from ancient marine deposits (Twisselmann 1967). Soils in areas of low relief are predominantly sandy loams, loams, and clay loams (Nelson et al. 1921, United States Department of Agriculture 1986, 1992). Rock outcrops are virtually nonexistent, although shallowly buried shale deposits occur in some Midway Valley sections (section = 2.6 km^2^).

The predominant shrub-steppe vegetation association is described as either the San Joaquin saltbush association (Küchler 1977) or Lower Sonoran grassland (Twisselman 1967). A dense ground cover of annual plants was dominated by Eurasian alien species, red brome (*Bromus rubens*) and red-stem filaree *(Erodium cicutarium).* Allscale (*Atriplex polycarpa*) was the most common shrub and was especially dense in disturbed areas along roadsides and edges of well pads. Other common shrubs included spiny saltbush (*Atriplex spinifera*), matchweed (*Gutierrezia bracteata*),California buckwheat (*Erigonoum fasciculatum*), cheesebush (*Hymenoclea salsola*), bladder pod (*Isomeria arborea)* and winterfat (*Krascheninnikovia lanata*). Areas adjacent to Buena Vista Lake playa were dominated by an alkali sink association characterized by inkweed (*Suaeda fruticosa*).

Petroleum was produced on both reserves beginning in 1919. However, production on NPR-1 remained low until 1977 when the Naval Petroleum Production Act of 1976 (Public Law 94-258) was implemented. Few areas on NPR-1 had well densities exceeding 25/km^2^. NPR-2 was intensively developed and much of it was considered to be depleted. Some portions had well densities exceeding 58/km^2^ but approximately 33% were older non-producing, abandoned, or suspended oil wells.

Sheep grazing disturbed much of the habitat on both reserves. Livestock grazing was proscribed on NPR-1 in the 1960s, but NPR-2 was still used as spring pasture for sheep. In areas where the flocks congregated around water trucks or bedded down, vegetation was eaten or trampled to bare mineral soil. Almost annually range fires consumed standing dead litter and shrubs over large acreages. There were few enforced restrictions on use of off-highway vehicles or hunting on NPR-2, but public access to NPR-1 was restricted.

### Transect Surveys

We slowly walked (< 2km/hr) straight line transects using hand-held sighting compasses and United States Geological Survey (USGS) topographic maps to maintain our bearings while we searched for kangaroo rat burrows within our fields of view. We walked two north-south transects through alternating columns of sections during a partial survey of NPR-1. Later we walked eight north-south transects spaced at 200-m intervals in all sections of both reserves. Potential giant kangaroo rat burrows had two or more of these characteristics: multiple horizontal entrances within an approximately circular mounded area of about 4 square meters of bare ground; vertical holes measuring about 5 cm in diameter that had no berms but were sometimes plugged with soft earth; and “haystacks” of clipped annual grass seed heads in the immediate vicinity of the mound. Potential burrows were plotted on USGS topographic maps. We live-trapped at selected burrows to confirm the presence of *D. ingens.* Detailed descriptions of survey methods, especially the information gathered to describe burrow systems, were published earlier (O'Farrell et al. 1987).

### Special Surveys

We selected five 65-ha survey sites in areas where giant kangaroo rat burrows were previously found in upland locations on NPR-1 (25R/36R, 3B/4B, 36R/1B, 28R, 32S)(sections and townships in the Mt. Diablo Meridian are identified by numeric/letter codes respectively). We walked sixteen line transects spaced at 50-m intervals through each site and gathered additional information on the systems within this unexpected habitat.

We also conducted an absolute count of burrows in Section 8B which contained what appeared to be optimal habitat based on historical descriptions. We located and characterized all burrows within five 1-ha belt transects within the southwest quarter of the southeast quarter of the section.

### Incidental Observations

Personnel conducting surveys for this and other wildlife studies visited both reserves on an almost daily basis. All giant kangaroo rat burrows that we located were documented and mapped.

### Soil Analyses

Because the soil series used by Grinnell (1932) were over 60 years old we contracted the United States Natural Resources Conservation Service (NRCS) to map them using contemporary descriptions which would more likely be used as updated soil surveys were completed throughout the remainder of the range of the species. Soil scientists observed the shape and steepness of slopes, drainage patterns, plant associations, and the types of parent material. They dug soil pits to study the characteristics of the soil profiles from the surface into the unconsolidated parent material. They used this information to develop models which enabled them to predict the soils of specific locations during mapping, and to assign taxonomic classes. The boundaries of the significant natural bodies of soil were drawn on aerial photographs obtained in 1984 and identified as a specific map unit and soil series. More detailed information on NRCS methodology can be found in their subsequent reports (United States Department of Agriculture 1986, 1992).

### Geographic Information System

Historical soils data were digitized into ARC/INFO from a 1:100,000 photo enlargement of Nelson et al.’s (1921) map. Recent data were digitized from 1:25,000 maps (United States Department of Agriculture 1986, 1992). Soils were digitized at the series level and aggregated, and area occupied by each series was measured and mapped. Locations of subsets of 272 and 290 giant kangaroo rat burrows located on NPR-1 and NPR-2, respectively, were plotted on 1:24,000 USGS topographic maps and digitized. Burrow locations were overlain with the soils maps to determine whether they were distributed within soil series in proportion to the area occupied by each series.

## Results

We observed a total of 1,080 giant kangaroo rat burrows on 30 sections of NPR-1 and five adjoining sections (5B, 6B, 17G, 19G, 31R), and on 22 sections of NPR-2 and one adjoining section (19B)(Fig. 2). Most were observed in clusters throughout Buena Vista Valley from sections between 31R and 18B eastward to Section 18G. Clusters of burrows were also found in Section 2D, NPR-2, and in sections 17S, 20S, 21S on the northeastern border of NPR-1. Isolated burrows were observed in an additional 19 sections of NPR-1 and 11 sections of NPR-2.

**Fig. 2.**
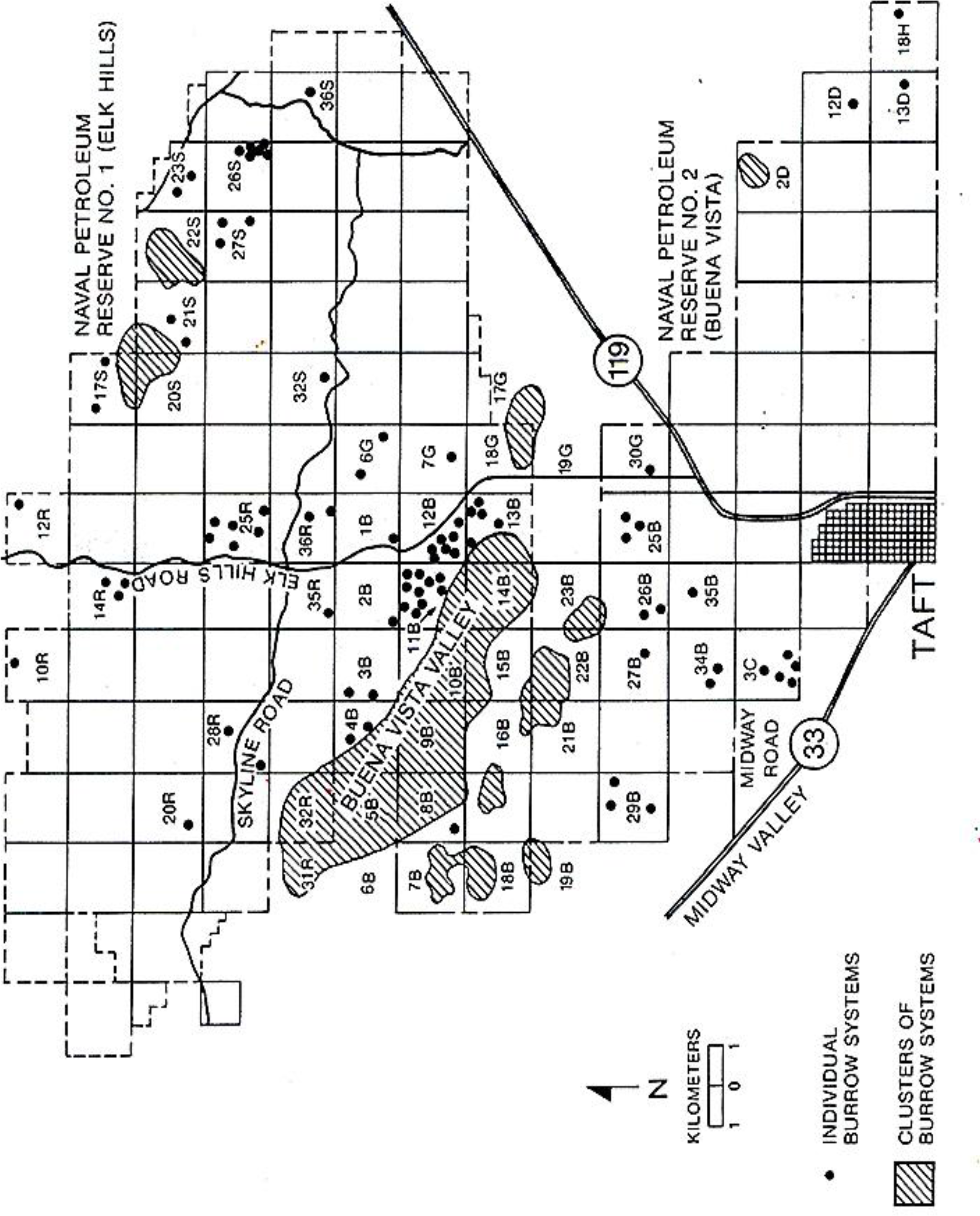
Locations where both clusters and isolated burrows of *D. ingens* were observed on the Naval Petroleum Reserves in California. Sections and townships in the Mt. Diablo Meridian are identified by numeric/letter codes respectively.

### Distribution Within Soil Types

Burrows were not distributed in proportion to the areal extent of soil series (Nelson et al. 1921) that were used by Grinnell (1932). On NPR-1 burrows were located in 4 of the 7 soils series, but 70% of the 272 mapped burrows were found in Panoche sandy loams that occurred in just 10% of the area (Table 1). Soils occupying 81% of the Reserve (Kettleman sandy loam, rough broken land) contained only 27% of the burrows (*χ* ^2^ = 1108, *d.f.* = 5, *P* < 0.001). On NPR-2 burrows were found on 6 of the 7 soil series and although Panoche loam and clay loam high phase occurred on just 2% of NPR-2, 50% of the 290 mapped burrows were observed in this series (Table 1). Soils distributed over 76% of NPR-2 (Kettleman sandy loams, Panoche sandy loams) contained 43% of the burrows (*χ* ^2^ = 3251, *df* = 6, *P* < 0.001).

**Table 1.**
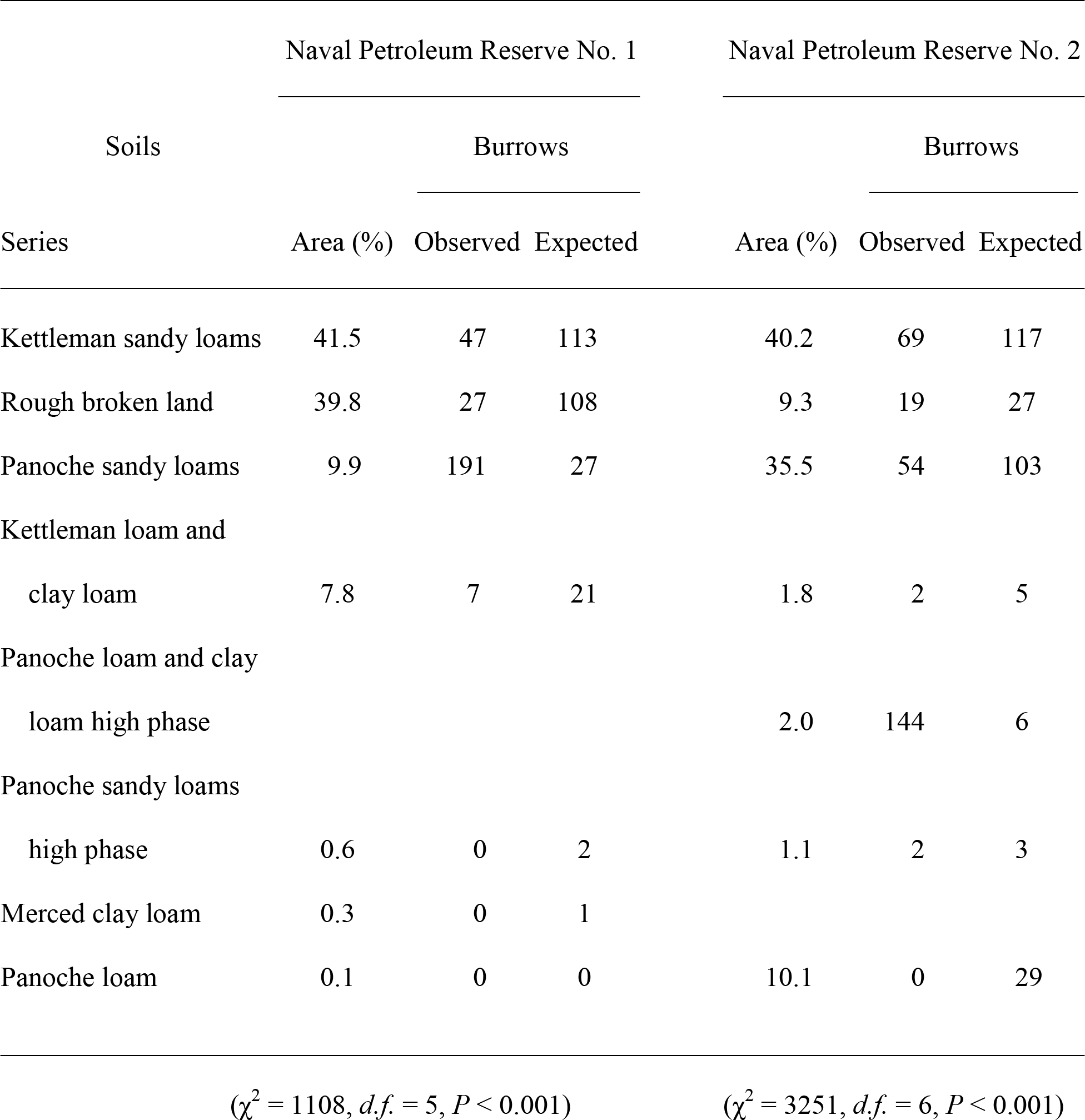
Expected and observed distributions of giant kangaroo rat burrows on Naval Petroleum Reserves in California based on proportions of areas overlain by various soil series described by Nelson et al. (1921). Results of chi-square tests indicate the significance of the disproportionate distributions.

The distribution of burrows within the revised soil series (United States Department of Agriculture 1986, 1992) showed a similar pattern. On NPR-1 burrows were located in 7 of 8 soil series, but 78% were observed in Kimberlina sandy loams that occupied only 14% of the area (Table 2). Soil series that occupied 77% of NPR-1 (torriorthents, Elk Hills sandy loam) contained only 19% of the burrows (*χ*^2^ = 935, *d.f.* = 7, *P* < 0.001). On NPR-2 burrows were located on 6 of 10 soil series, but 74% were observed in Kimberlina sandy loam and Tupman gravelly sandy loams that occupied 28% of the Reserve (Table 2). Soil series occupying 56% of NPR-2 (torriorthents, Guijarral sandy loam, Elk Hills sandy loam) contained 24% of the burrows (*χ* ^2^ = 347, *df* = 9, *P* < 0.001).

**Table 2.** Expected and observed distributions of giant kangaroo rat burrows on Naval Petroleum Reserves in California based on proportions of areas overlain by various soil series (U.S. Department of Agriculture 1986, 1992). Results of chi-square tests indicate the significance of the disproportionate distributions.

| Soils<br>Series | Naval Petroleum Reserve No. 1 |  |  | Naval Petroleum Reserve No. 2 |  |  |
| --- | --- | --- | --- | --- | --- | --- |
|  | Area (%) | Burrows |  | Area (%) | Burrows |  |
|  |  | Observed | Expected |  | Observed | Expected |
| Torriorthents | 55.7 | 28 | 152 | 28.3 | 24 | 82 |
| Kimberlina sandy loam | 14.0 | 212 | 38 | 24.5 | 177 | 71 |
| Elk Hills sandy loam | 20.9 | 23 | 57 | 13.0 | 26 | 38 |
| Guijarral sandy loam |  |  |  | 14.6 | 20 | 42 |
| Elk Hills complex | 5.1 | 1 | 14 | 2.0 | 0 | 6 |
| Tupman gravelly sandy<br>loam |  |  |  | 3.2 | 38 | 9 |
| Kimberlina-Cajon<br>riverwash | 1.7 | 6 | 5 | 0.1 | 0 | <1 |
| Kimberlina-urban land | 0.9 | 1 | 2 | 2.7 | 0 | 8 |
| Cajon sandy loam | 0.3 | 1 | 1 | 1.3 | 0 | 4 |
| Others combined | 1.3 | 0 | 4 | 10.2 | 5 | 30 |

( $\chi^2 = 935$ , *d.f.* = 7, $P < 0.001$ )
( $\chi^2 = 347$ , *d.f.* = 9, $P < 0.001$ )

### Position on Slope and Slope Angle

Of a subset of 170 burrows found on NPR-1, 63% were found on the floor of the valley, 23% on the crests of ridges, 5% on upper slopes, 1% on middle slopes, and 1% on lower slopes. Regardless of their position on slopes burrows were found most consistently in areas of low relief. Slope angles of 32 burrows ranged between 1-17° and averaged 4.3 ± 0.5° in the valleys. Slope angles of 39 burrows in the uplands ranged between 2-23° and averaged 6.4 ± 0.8°. Averdage slope angles were significantly different between the two slope positions (t = 2.21, df = 69, P < 0.05).

### Proximity to Human Disturbances

The distribution of giant kangaroo rats on the reserves overlapped areas of little to intense petroleum development. The highest densities of burrows were in the valley which was virtually devoid of petroleum developments. However, isolated burrows scattered in the hills were in close proximity to various disturbances. Well pads were the most frequently recorded (73%) of 211 human disturbances in the vicinity of 71 burrows, but on average they were the most distant (150 ± 61 m, n=150). Pipelines were the closest disturbances (38 ± 11 m, n=8) but were observed least frequently (4%). Dirt roads were observed within an average distance of 50 ± 10 m (n=28) from 13% of the burrows. Miscellaneous disturbances such as power lines, firebreaks, graded areas, tank farms, fences, and sheep grazing, were observed an average of 52 ± 12 m (n=21) from 10% of the burrows.

## Discussion

Giant kangaroo rat burrows were widely distributed, and, in some areas, were relatively abundant on both reserves. The greatest densities were found in Buena Vista Valley on a gently sloping alluvial plain that supported a dense ground cover of winter annuals and a relatively sparse shrub cover. Previous studies indicated that preferred habitat was characterized by open, flat to gently rolling terrain lacking sharp physiographic features (Grinnell 1932; Shaw 1934; Hawbecker 1951, Williams 1992, Williams et al. 1995), Vegetation in such habitat generally consisted of a dense ground cover of grasses and forbs with no shrub cover (Shaw 1934; Hawbecker 1944, 1951). Portions of Buena Vista Valley, especially Section 8B, contained what was historically described as optimal habitat for the species.

Most burrows were found in low relief habitats on the valley floors adjacent to NPR-1, but 28% were found on upper slopes or the crests of ridges. The number and wide distribution of burrows in the upland terrain was unexpected. Previous reports indicated that giant kangaroo rats were virtually absent from areas of high relief (Grinnell 1932; Shaw 1934; Hawbecker 1944, 1951). It appeared that suitable giant kangaroo rat habitat occurred in the uplands as small widely scattered islands on ridges or crests of slopes wherever slope angles were reduced below 10° and soils were deep and stone-less. Concurrently we discovered a colony of approximately 20 active giant kangaroo rat burrow systems on Monocline Ridge in the Ciervo Mountains 175 km northwest of the reserves (P. M. McCue, personal communication). They were located at an elevation of 613 m (436 m above the San Joaquin Valley floor) on a plateau that faced 337° and had a slope angle of 5-9°.

Our results using the older soil series classification (Nelson et al. 1921) largely confirmed Grinnell's (1932) hypothesis that the distribution of giant kangaroo rat habitat appeared to nearly coincide with the occurrence of Panoche fine sandy loam. Panoche fine sandy loam was characterized by loose calcareous surface soils that were extremely friable and extended to depths of 36-46 cm; subsurface soils were loose, calcareous and permeable, and extended to 76-102 cm (Cole et al. 1945). Burrows were not found in alkaline soils in valley bottoms. Grinnell (1932) suggested that such areas were avoided because they were periodically flooded.

The preference for specific soil series remained when the areal extent of the revised soil series nomenclature (United States Department of Agriculture 1986, 1992) was used. The greatest densities of burrows coincided with the occurrence of Kimberlina sandy loam and Tupman gravelly sandy loam. Both are very deep (115-150 cm), well drained soils formed in alluvium derived from granitic and sedimentary rock sources. Kimberlina sandy loam was formed on recent alluvial fans having slopes of 0-2° and is virtually stone-free. Tupman gravelly sandy loam was formed on flood plains and fans having slopes between 0-15° and contains 10-15% clay and 10-30% rock fragments (United States Department of Agriculture 1986, 1992).

Grinnell (1932) observed that the texture of the soil where burrows were found was,“…marvelously suited to year-long maintenance of burrows—firm, holding together well, yet ‘diggable’ for a rodent of weak fossorial power,” and that it had,“…no stones to strike, no resistant ‘adobe;’ yet no tendency to cave in.…” He speculated (Grinnell 1932) that the species “…finds optimum conditions for burrowing in a belt of country where the rainfall (5 inches or less per annum is not sufficient to wet the ground thoroughly…to a depth greater than the animals can easily exceed in their diggings.” He concluded that, “The prime basis for the presence of this particular rodent would thus be a rainfall within some certain annual maximum, plus a particular kind of soil as regards permeability by water and diggability by the animal”

Williams (1992) located occupied habitats in a wide variety of soil types, but the largest extant colonies were associated with loam series soils. In the northern part of the species’ range (western Fresno and eastern San Benito counties) 79 colonies were found on 12 soil series, but 66% were located on just 4 series (Williams et al. 1995). They concluded that the largest colonies, that included approximately 93% of the estimated population, were located on 3 of the soil series. They did not provide information on the areal extent of the soil series which precludes testing whether colonies were distributed preferentially.

Results of this study were incorporated into DOE’s Biological Assessment used to successfully complete an ESA Section 7 Consultation with FWS regarding the potential effects of petroleum production activities on the giant kangaroo rat. An important conservation measure was to include the species in the pre-activity survey process (Kato and O’Farrell 1987) to mitigate possible negative impacts. The guidelines emphasized that, “ Direct impacts to giant kangaroo rat burrow systems must be avoided.” Minimal 10-meter buffers were required around each burrow system to minimize the potential for accidental damage, and larger buffers were prescribed in cases where the extra distance was required to ensure the physical integrity of the burrow systems.

Giant kangaroo rat burrows occurred most frequently, and in the greatest densities, on the valley floors that were not underlain by extensive petroleum deposits, and where the potential for negative impacts was low. The majority of development took place on the dissected uplands that were underlain by rich oil deposits. Burrows in these areas were potentially vulnerable to disturbances or destruction, but little evidence of such damage was observed. During 296 pre-activity surveys conducted over a 4-year span, only 8 burrows were found in the vicinity of proposed construction projects covering approximately 14.5 km^2^ (Kato et al. 1985). No projects had to be modified to avoid negative effects on the species.

FWS is responsible for implementing ESA and conducts species status surveys, prepares listing documents for worthy species, defines critical habitats, issues biological opinions for federal projects, designs and implements recovery plans, and reviews conservation plans for Section 10 non-federal actions that require an incidental take permit. Our results should help FWS conserve and ultimately recover the species.

When the giant kangaroo rat was listed as endangered in 1987 FWS chose not to designate critical habitat. Our confirmation of the correlation between optimal habitat and contemporary soil series could be used as a discriminator for helping to define critical habitat, focus recovery actions (United States Fish and Wildlife Service 1998) in appropriate soil series, and evaluate potential impacts of actions proposed in preferred soil types.

We also discovered that the distribution of giant kangaroo rats was not limited to valley floors, and that burrow systems were widely distributed in suitable upland terrain. The data suggest that species status surveys should not be limited to lowlands based on earlier descriptions of preferred habitat. If the species is distributed in other suitable uplands off of the reserves, as we discovered in the Ciervo Hills, the species may be more widespread and abundant than previously thought.

Our results also show that large scale petroleum developments can take place within the range of the giant kangaroo rat if suitable mitigation measures, including pre-activity surveys, are designed and implemented. Implementing pre-activity surveys helped protect burrow systems, but did not interfere with completion of major projects in a timely way, nor did it add significant costs. On NPR-1 DOE achieved peak production in excess of 200,000 barrels of petroleum per day while concurrently conserving this endangered species.

## ACKNOWLEDGMENTS

This study was supported by the United States Department of Energy, Office of Naval Petroleum and Oil Shale Reserves, and Chevron U.S.A. Inc., through the Nevada Operations Office under Contract No. DE-AC08-93NV11265. We are grateful for the assistance provided by our colleagues who helped gather some of the information synthesized here: W. H. Berry, B. G. Evans, D. B. Hardenbrook, C. E. Harris, R. Horwitz, J. W. Johnson, T. T. Kato, D. Land, J. S. McManus, P. C. Muick, G. Orth, D. Padley, M. Spencer, and M. Whitsel. D. Kliman of EG&G/EM’s Remote Sensing Laboratory digitized and entered information into the geographic information system and provided analyses of the distributions of soil series and burrow locations. R. L. Norland, formerly of Williams Brothers Engineering Co. which operated NPR-1, helped obtain the funding. D. F. Williams of California State University, Stanislaus, shared his knowledge of the species and its burrows with us. R. J. Hensel provided helpful editorial comments. M. M. Shaw translated the abstract into Spanish. Permission to live-trap and handle this species prior to its Federal listing was granted in a Memorandum of Understanding from the California Department of Fish and Game.

